# A Methodological Assessment and Characterization of Genetically-Driven Variation in Three Human Phosphoproteomes

**DOI:** 10.1101/271650

**Authors:** Brett W. Engelmann, Chiaowen Joyce Hsiao, John D. Blischak, Yannick Fourne, Michael Ford, Yoav Gilad

**Affiliations:** Department of Human Genetics, University of Chicago, Chicago, Illinois, USA; Department of Medicine, University of Chicago, Chicago, Illinois, USA; MS Bioworks, LLC, 3950 Varsity Drive, Ann Arbor, Michigan, USA

## Abstract

Phosphorylation of proteins on serine, threonine, and tyrosine residues is a ubiquitous post-translational modification that plays a key part of essentially every cell signaling process. It is reasonable to assume that inter-individual variation in protein phosphorylation may underlie phenotypic differences, as has been observed for practically any other molecular regulatory phenotype. However, we do not know much about the extent of inter-individual variation in phosphorylation because it is quite challenging to perform a quantitative high throughput study to assess inter-individual variation in any post-translational modification. To test our ability to address this challenge with current technology, we quantified phosphorylation levels for three genotyped human cell lines within a nested experimental framework, and found that genetic background is the primary determinant of phosphoproteome variation. We uncovered multiple functional, biophysical, and genetic associations with germline driven phosphopeptide variation. Variants affecting protein levels or structure were among these associations, with the latter presenting, on average, a stronger effect. Interestingly, we found evidence that is consistent with a phosphopeptide variability buffering effect endowed from properties enriched within longer proteins. Because the small sample size in this ‘pilot’ study may limit the applicability of our genetic observations, we also undertook a thorough technical assessment of our experimental workflow to aid further efforts. Taken together, these results provide the foundation for future work to characterize inter-individual variation in post-translational modification levels and reveal novel insights into the nature of inter-individual variation in phosphorylation.

## INTRODUCTION

Protein phosphorylation is a ubiquitous mediator of information flow in essentially all cellular processes [1–4], with a recent survey estimating that roughly 75% of the proteome can be phosphorylated [5]. Dysregulation of protein phosphorylation has long been recognized as a driver of disease [4, 6–8], and plays an important role in achieving and maintaining every ‘hallmark’ of cancer [9]. While the proteins involved and mechanistic details of the major phosphorylation mediated signal transduction pathways are largely known [2], a growing body of research seeks to understand phosphorylation mediated information transfer as an integrated system using broad, quantitative, and unbiased surveys of the phosphoproteome combined with other ‘omic’ data [10–13]. Recent advances in liquid chromatography coupled to tandem mass spectrometry (LC-MS/MS) technology have enabled such surveys [5, 14, 15], and multiple
studies have reported the analysis of LC-MS/MS phosphoproteomic data together with genomic, transcriptomic, proteomic and metabolomic data [5, 16–21].

In particular, integrative phosphoproteomic-genomic studies have provided further evidence of the importance of phosphorylation in evolution and disease. Previous studies have combined genomic data with phosphoproteomic data to provide evidence that phosphorylation sites are conserved across species [22, 23], are under evolutionary constraint in humans [24], and are over-represented in mutations that cause diseases in humans [24, 25]. Phosphoproteomic data has also been combined with genomic and protein-binding specificity data to develop models that predict mutations likely to alter phosphorylation signaling in cancer [26] or perturb specific kinases [27]. More recently, integrative phosphoproteomic-genomic studies have improved our understanding of how genetic alterations impact phosphorylation mediated signaling by combining LC-MS/MS derived quantitative phosphoproteomic and genomic data from the same samples. A recent integrative study identified signaling pathways that are differentially activated in breast cancer samples depending upon the mutation pattern of a frequently mutated gene [19]. In another example, phosphoproteomic data and exome sequence data collected from multiple ovarian cancer cell lines was used to assess the impact a subset of genetic variants have on a predicted phosphoprotein network state [28]. Despite this progress, we are not aware of any studies that have systematically characterized how genetic variation affects variation in phosphorylation levels across a set of commonly measured samples. Moreover, because many of the preceding *in vivo* studies were performed on cancer models, the contribution of heritable variation to naturally occurring inter-individual differences in protein phosphorylation levels remains unexplored.

Quantitative trait locus (QTL) mapping is a powerful approach to analyze inter-individual variation in phosphorylation levels. When QTL mapping is applied to molecular phenotypes, such as mRNA or protein expression levels, these are treated as quantitative traits. The goal of regulatory QTL mapping is to identify associations between inter-individual variation in the molecular phenotypes and the corresponding genotypes from multiple individuals [29]. Recent progress cataloging QTLs associated with various molecular phenotypes using high throughput approaches has been rapid [30–40]. Yet, to date, there have been no quantitative studies with an aim to characterize inter-individual variation in post-translational modification (PTM) levels. To begin addressing this gap, we performed a pilot study to assess the feasibility of QTL mapping of PTM levels. We applied liquid chromatography coupled to tandem mass spectrometry (LC-MS/MS) to derive quantitative phosphoproteomes from three HapMap [41] lymphoblastoid cell lines (LCLs) donated from Yoruba (Ibadan, Nigeria) individuals. Along with genomic information [42], other quantitative datasets, including transcriptomic [31] and proteomic [34], have been previously collected from these LCLs. We leveraged these previous data sets and the quality of our phosphoproteomic data to explicitly estimate phosphopeptide variance arising from the genetic background. We found that the genetic background drives the majority of the observed variance, and uncovered many novel relationships between germline genetically-driven phosphorylation variation and diverse molecular annotations. We also included a power analysis with varying levels of increasing technical variance to aid the design of future studies.

## RESULTS

### Nested deep quantitative phosphoproteome profiling

We applied a nested experimental design in order to characterize variation in protein phosphorylation between samples. We aimed to estimate the relative contributions from biological and technical sources to the observed variance in phosphopeptide quantification. We designed the study to specifically allow us to consider the contributions of genetic background, tissue culturing, and MS processing (**Figure 1**). We employed Stable Isotope Labeling by Amino Acids in Cell culture (SILAC) [43, 44] for relative quantitative comparisons of phosphopeptides using a common unlabeled reference LCL, and labeled sample LCLs (**Supplemental Figure 1**). The phosphoproteome data set contains 192 1.5 hr gradient LC-MS/MS experiments on a Q-Exactive quadrupole orbitrap [45], employing higher energy collisional dissociation to fragment peptides. Using this experimental approach, combined with the MaxQuant [46] proteomic software suite and Andromeda [47] search engine, we identified over 22,000 phosphopeptides from 5,143 unique protein groups at an FDR of 1% (**Table 1, Supplemental Table 1, Supplemental Figure 2**). Ultimately, 17,774 phosphopeptides mapping to 4,584 protein groups produced spectra enabling confident localization of the site of phosphorylation and were assigned to SILAC pairs (‘Class 1’ quantifications, **Table 1**).

**Figure 1.**
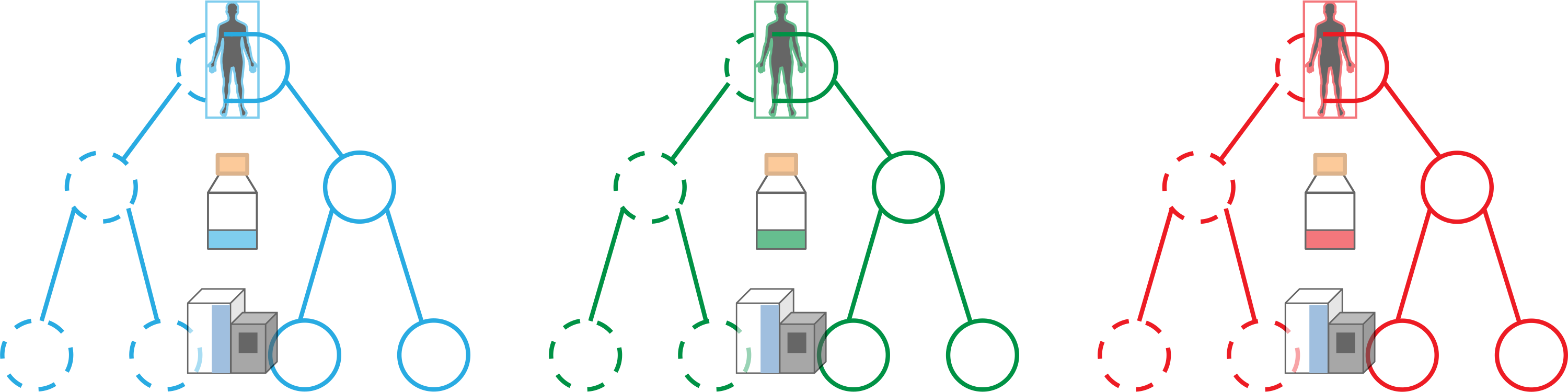
Nested experimental design. HapMap LCLs derived from three Yoruba males in Ibadan, Nigeria were repeatedly cultured and repeatedly subjected to a multistep mass spectrometry workflow (see Supplementary Figure 1). Dotted vs un-dotted circles represent the two different processing batches.

**Table 1.** Phosphopeptide level MS summary. All sites identified at an FDR of 1%

| Identified |  | Quantified |  | Class 1† quantified |  | Class 1† quantified in each* LCL |  | Class 1† quantified in each* culture replicate |  | Class 1† quantified and protein normalized in each* culture replicate |  |
| --- | --- | --- | --- | --- | --- | --- | --- | --- | --- | --- | --- |
| Phospho peptides | Proteins | Phospho peptides | Proteins | Phospho peptides | Proteins | Phospho peptides | Proteins | Phospho peptides | Proteins | Phospho peptides | Proteins |
| 22766 | 5143 | 21944 | 4845 | 17774 | 4584 | 11117 | 3514 | 4742 | 2073 | 3257 | 1181 |

†Refers to subset of phosphorylation sites with median localization probability of .99 and min .75
\*Refers to the intersection

### Donor identity is the main biological source of phosphoproteome variation

As a first step of our analysis, we used normalized values (median-adjusted and quantile-normalized, see methods) to examine phosphopeptide variation prior to accounting for variation in protein expression levels. We applied principal component analysis (PCA) to this dataset and found that PC1 was associated with processing date, and PC2 was associated with donor identity (**Supplemental Figure 3A**). These results indicated that a processing date batch effect is associated with substantial technical variation in our measurements. Thus, we applied the empirical Bayes approach ComBat [48] to estimate and regress this batch effect from the data, applied PCA to the residuals, and visually determined that data across samples cluster by donor individual (**Supplemental Figure 3B**). Following these results, the batch-corrected, normalized values were applied throughout our analysis.

To explicitly account for the confounding effect of variation in protein expression levels, we assigned relative protein levels to each phosphopeptide using SILAC ratios derived from a previously reported MS dataset collected from 60 Yoruba LCLs, which employed the same reference sample [34, 49]. We processed these data with MaxQuant, yielding 3,885 identified and quantified protein groups in each of the three LCLs (the intersection) that we used in the current study (at a peptide and protein FDR of 1%; see methods, **Supplemental Table 2**). Because these SDS-PAGE protein expression levels were derived separately from the phosphopeptide data, we had to perform a separate normalization step and batch effect correction. We thus adopted a strategy that is commonly used in regulatory QTL studies to maximize the accuracy of molecular measurements. Specifically, to account for noise within the protein expression data we leveraged the available genotypes of the 60 LCLs to detect protein QTLs. Given that power to detect QTLs depends on the accuracy of the measurements, and that genotype distributions across *cis* regulatory loci are mostly uncorrelated, it is reasonable to assume that the protein data matrix that produces the most protein QTLs contains the most accurate protein estimates (indeed, an assumption shared by most regulatory QTL studies [31, 34, 50, 51]). In order to identify this matrix, we iteratively applied PCA to the protein data and regressed unidentified confounders to maximize the number of protein QTLs identified across all 60 LCLs (see methods). Following the empirical correction of noise within the protein expression data, 1,181 protein groups were assigned to 3,257 phosphopeptides present in each of the three LCLs subjected to the phosphoproteomic work-up (**Table 1**).

We used the corrected protein expression levels and the batch-effect corrected phosphopeptide values to estimate contributions to variance from the genetic background (the donor), culture replication (technical replication of the cell culture) and technical workup (protein sample processing and MS workflow). We fit a nested random effects model to each phosphopeptide with corrected protein expression levels as a covariate (see methods). We found that, for both absolute and relative phosphopeptide variance distributions, the genetic background dominates the observed variance (**Figure 2A and 2B**). We also fit our nested random effects model with normalized phosphopeptide measurements in order to assess the impact our batch-effect correction approach had on our estimates. After including both processing batch and corrected protein levels as fixed-effect covariates, we found that the dominant contribution of the genetic background to the observed variance did not change (**Supplemental Figure 4**).

**Figure 2.**
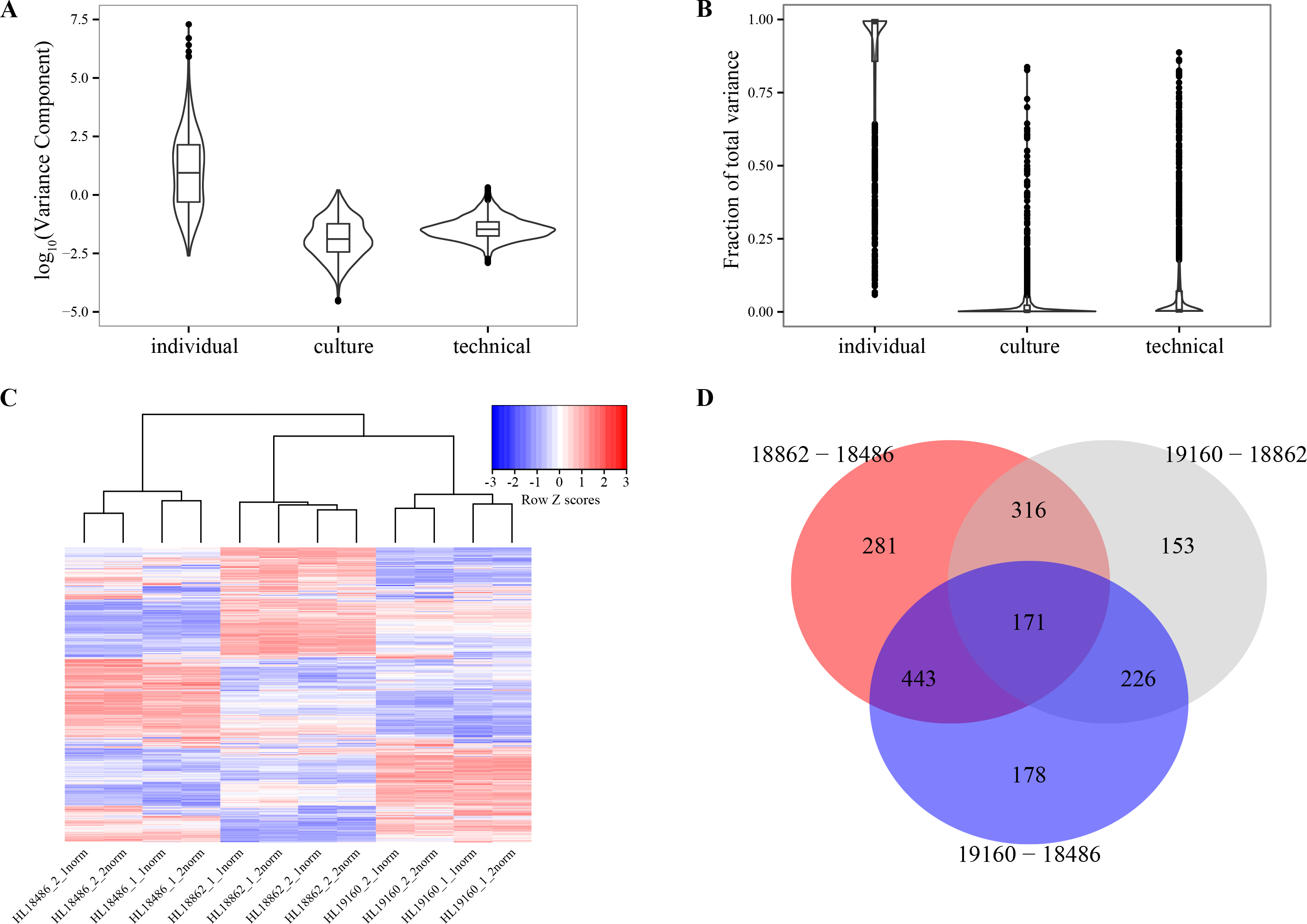
Genetically-driven phosphoproteomic variation. Violin plots of **(A)** absolute and **(B)** standardized phosphopeptide variance components derived from each layer of the hierarchical design after accounting for protein levels. **(C)** Heatmap of protein normalized phosphopeptide SILAC ratios. (**D)** Venn diagram of differential phosphorylation results across all three pairwise inter-individual comparisons (FDR 5%).

Next we considered differences in phosphorylation levels across the three LCLs. The study was not designed with a main aim to provide mechanistic insight into the specific pathways that drive inter-individual variation across LCL phosphoproteomes. Indeed, there is no specific stimulation response of interest and this is a small sample with which to attempt such analysis, just three individuals. Nevertheless, we hierarchically clustered the batch effect corrected and protein expression normalized phosphopeptide SILAC ratios and again found that the data cluster by donor (**Figure 2C**). To focus our analysis on phosphorylation levels, we modeled the batch effect corrected phosphopeptide data while accounting for protein expression levels (see methods). Using this approach, we classified 48% (1577 of 3257) of the phosphopeptides as differentially phosphorylated between individuals (omnibus F test; FDR of 5%). We observed modest effect sizes (**Supplemental Figure 5**) and a varied complement of differentially phosphorylated peptides and enriched gene ontology categories across inter-individual comparisons (**Figure 2D, Supplemental Tables 3-5**). We also observed a variety of differentially phosphorylated phosphopeptide sequences, with only two kinase motifs enriched at an FDR of 5% (**Supplemental Table 6**) and no single amino acid enriched at any sequence position across any inter-individual comparison (**Supplemental Figure 6**). These finding demonstrate that there is an extensive amount of modestly varying phosphorylation across three non-stimulated LCLs derived from genetically different, albeit closely related donors.

### Characterization of genetically-driven differential phosphorylation

Understanding the nature of the genetic differences that putatively drive variation in protein phosphorylation between individuals is of fundamental interest and aids further experimentation. A key to this understanding is an assessment of the impact variants mapping to different functional categories have on inter-individual phosphopeptide variation. We undertook this assessment with an enrichment analysis.

To begin our assessment, we used the genomic sequence information available from all three donors [42]. Genetic variants impact phosphopeptide levels by altering protein expression or function. The former were previously captured within this system as protein QTLs (pQTLs) [34], while the latter manifest via amino acid coding variants (non-synonymous SNPs and indels). We investigated all identified genetic differences between the three individuals in our study. In this case, though our sample size is small, the analysis relies on the large number of phosphopeptides we measured, and thus is not as underpowered as it may intuitively seem. Specifically, we are able to consider 19,002 coding variants affecting 8,656 unique genes and 181 pQTLs in these three individuals.

Using the p-values derived from the inter-individual F-tests we described above, we calculated (in a threshold independent fashion) the Spearman’s correlation between various genetic annotations and phosphopeptide variation [34] (**Table 2**, see methods). A significant positive correlation between the presence of an annotation and phosphopeptide variability is indicative of an ‘enrichment’ of that annotation amongst proteins that contain phosphopeptides that are highly variable between individuals. Given that the expression level of a protein may impact phosphorylation levels in *cis* through enzyme-substrate titration, we hypothesized that proteins with annotated pQTLs would be enriched amongst those proteins that contain highly variable phosphopeptides. Consistent with this, we observed a significant enrichment of proteins with a pQTL (p = 0.03, **Table 2**). We also hypothesized that coding variation within a protein would correlate positively with phosphopeptide variability (in *cis*). Indeed, we found that proteins containing at least one non-synonymous variant were enriched (strongly, relative to proteins with annotated pQTLs) amongst those proteins that contain highly variable phosphopeptides (p = 6.48 × 10^−8^, **Table 2**).

**Table 2.** SNP categorical enrichment analysis

| Annotation (Protein) | N (phosphopeptides) | Background | p-value |
| --- | --- | --- | --- |
| At least one coding variant | 3257 | Phosphopeptides subjected to F-Test | $6.48 \times 10^{-8}$ |
| pQTL | 3257 | Phosphopeptides subjected to F-Test | $2.67 \times 10^{-2}$ |
| At least one variant within a Pfam domain | 1332 | Phosphopeptides within proteins that have at least one coding variant | $9.83 \times 10^{-6}$ |
| At least one variant within a disordered region | 1332 | Phosphopeptides within proteins that have at least one coding variant | $7.51 \times 10^{-1}$ |
| At least one PolyPhen HVAR Deleterious variant | 1332 | Phosphopeptides within proteins that have at least one coding variant | $6.08 \times 10^{-6}$ |
| At least one PolyPhen HDIV Deleterious variant | 1332 | Phosphopeptides within proteins that have at least one coding variant | $1.56 \times 10^{-3}$ |
| Phosphorylation regulation | 1332 | Phosphopeptides within proteins that have at least one coding variant | $8.22 \times 10^{-1}$ |
| At least one variant within phosphorylation regulation domain | 273 | Phosphopeptides within proteins that have at least one coding variant in a Pfam domain | $5.75 \times 10^{-1}$ |

As a control for our approach we also tested whether the number of coding variants within a protein is correlated with inter-individual variability in phosphorylation levels. This should not be the case because the functional and biophysical context of a variant within a protein should have a greater impact on phosphopeptide variation than the overall number of variants within a protein. We tested this hypothesis by limiting the background set of phosphopeptides analyzed to those that are within proteins containing nonsynonymous variants. Within this background, we found that proteins with multiple coding variants are indeed not significantly enriched amongst those proteins that contain more variable phosphopeptides (>1 coding variant; n=1,332, p=0.11).

Thus, we proceeded by investigating the relationship between the context of genetic variants and phosphopeptide variation. We considered variant placement within the dichotomy of structured globular domains or disordered protein segments. While protein domains are the modules that largely impart protein function [52, 53], disordered regions contain most of the phosphopeptides observed to date [54] (here 73%) and play critical roles in signal transduction and macromolecular assembly [55–57]. To supplement this dichotomy, we also categorized variants as likely or unlikely to impact protein function using PolyPhen-2 [58], which is an empirically trained prediction algorithm that considers multiple sequence and structure features. We limited the background set of phosphopeptides analyzed to those that are within proteins containing at least one nonsynonymous variant. Within this background, we found that both proteins with variants mapping to defined units of protein structure (domains) and proteins with variants likely to impact function (PolyPhen-2 [58] “deleterious” variants) are enriched amongst those that contain highly variable phosphopeptides (all p-values <= 1.56 × 10^−3^; **Table 2**). However, we found that proteins with variants mapping to disordered regions are neither enriched nor depleted among those that contain highly variable phosphopeptides (p = 0.751, **Table 2**).

The function of a protein may also influence the likelihood that a variant impacts phosphorylation levels. Given their centrality within phosphorylation signaling networks, we investigated if proteins that regulate phosphorylation signaling are particularly impacted by genetic variation. Phosphorylation regulation proteins (PRPs) – kinases, phosphatases, and proteins containing non-catalytic phosphopeptide-recognition domains – alter phosphorylation levels in *cis* directly via catalysis or indirectly via spatial organization and subsequent catalysis by interacting proteins [1, 59, 60]. Notably, we did not find an enrichment of PRPs amongst proteins that contain highly variable phosphopeptides (nonsynonymous variant background, p = 0.82, **Table 2**). While a mutation within a PRP does not increase phosphopeptide variation relative to other proteins with mutations, it is possible that a targeted mutation within a kinase, phosphatase, or phosphopeptide-recognition domain is more likely to predispose such proteins toward increased phosphopeptide variability compared to mutations impacting other domains. To investigate this possibility, we limited the background set of phosphopeptides analyzed to those mapping to proteins that contain at least one variant in a domain. Within this background, we again did not find an enrichment of proteins with mutations in phosphopeptide regulation domains (p = 0.57, **Table 2**). Put together, these results indicate that the context of a mutation within a protein is the primary determinant of its ability to impact phosphopeptide levels in *cis*, regardless of any association that protein may have to phosphorylation regulation.

Lastly, we investigated the contextual impact variants may have on peptide-motif mediated interactions. Motif mediated protein-protein interactions and catalysis are important signal transduction mechanisms [61]. Binding of short linear peptides and phosphopeptides by PRPs directs specific catalysis and the formation of transient protein-protein interactions during signal transduction [1, 61]. Motif defining amino acids are typically found within +/− 5 residues of the phosphorylation site [62–66], with other residues extending beyond the motif also playing an important role to ensure interaction specificity [67]. While we were not able to obtain a large enough sampling of variants that disrupt annotated motifs (here only 5) to perform enrichment analysis, we did observe increased phosphopeptide variability as the distance between the phosphorylated site and the closest variant (in *cis*) decreased (R = −0.11; p = 4.37 × 10^−5^, **Supplemental Figure 7**). This proximity effect is consistent with a signature of altered motif mediated protein-protein interactions.

### The putative functional impact of differences in phosphorylation

In order to further our understanding of differential phosphorylation we also analyzed the characteristics of the proteins and phosphopeptides associated with phosphorylation variation without explicit regard to coding variation. The characteristics we uncovered may not be specific to phosphopeptide variation driven by genetic differences, but generalizable to phosphopeptide differences driven by *eg* drug treatment. To do this, we carried out the same enrichment analysis approach outlined above but employed phosphopeptide or protein, rather than genetic, annotations.

Phosphorylation events may or may not result in changes to protein function [68, 69]. Indeed, while phosphorylated sites are more conserved than non-phosphorylated sites [8, 70, 71], this conservation is greatly increased when only considering phosphopeptides that have a known function [68, 72]. Following batch-effect correction, the majority of the phosphopeptide variance observed in our study is derived from the genetic background rather than noise sources (**Figure 2A and 2B**). Therefore, we hypothesized that phosphopeptides that map to regions of annotated function would be enriched amongst highly variable phosphopeptides. Indeed, we observed an enrichment of phosphopeptides that map to functional protein segments (domains or annotated motifs) amongst highly variable phosphopeptides (p = 6.50 × 10^−4^ (domains); p = 2.06 × 10^−7^ (motifs), **Supplemental Table 7**). Next, we asked whether phosphopeptides that map to phosphopeptide-regulation domains have altered variability relative to phosphopeptides that map to other domains. To test this, we limited the background set of phosphopeptides to those within domains. Within this background, we did not find an enrichment of phosphopeptides that map to phosphopeptide-regulation domains amongst highly variable phosphopeptides (p = 0.28, **Supplemental Table 7**). We also found no relationship between the functional association of a protein to phosphopeptide regulation and phosphopeptide variability (p = 0.06, n = 3257). These findings again support the notion that PRPs do not possess more variable phosphosites relative to other proteins and the context of a phosphopeptide within a protein is the primary indicator of its penchant for variability.

As noted above, phosphorylation sites are predominantly found within disordered regions between domains. Yet, we observed a depletion of phosphosites that reside within disordered segments amongst highly variable phosphopeptides (p = 6.00 × 10^−3^, **Supplemental Table 7**). This result was somewhat unexpected given that disordered regions are enriched in functional motifs [55, 73] and our observation above that phosphopeptides mapping to annotated motifs display (on average) increased variability relative to all phosphopeptides (**Supplemental Table 7**). We therefore hypothesized that the functional properties of proteins at the systems level may contribute to this observation. The relative importance of proteins within an interaction network may impact phosphopeptide variability in *cis*. For example, proteins with a high degree of connectivity (hubs) are more likely than proteins with a low degree of connectivity to be essential [74]. Protein hubs tend to be long, highly modified, and enriched in regions of structural disorder [75, 76].

Using externally derived annotations (see methods), we found that proteins with more interactions, more PTMs, and higher disordered residue content (captured by the percentage of disordered residues and the longest run-length of disordered residues within a protein) are depleted amongst those with highly variable phosphopeptides (all p-values <= 0.03; **Supplemental Table 8**). Consistent with these annotation-derived observations, we also observed a depletion of highly phosphorylated proteins amongst those with highly variable phosphopeptides (p = 4.27 × 10^−6^, **Supplemental Table 8**). These observations could be driven in part by multiple mechanisms to direct specific protein-protein interactions such as the coordination between multiple PTMs, domains, and linear motifs that may be more common for longer proteins [3, 55, 56, 77, 78]. Indeed, we found longer proteins are depleted amongst those with highly variable phosphopeptides (p = 2.04 × 10^−5^, **Supplemental Table 8**).

The generally lower expression levels of longer proteins may also contribute to more “robust” phosphopeptide signaling due to mass action effects [79, 80]. According to this hypothesis, lowly expressed proteins are less susceptible to PTMs resulting from promiscuous moderate affinity interactions. Consistent with this, we found lowly expressed proteins are depleted amongst those with highly variable phosphopeptides (p = 5.88 × 10^−6^, **Supplemental Table 8**). Taken together, these results indicate that the systems-level properties of a protein significantly impact the likelihood that phosphorylation levels will be altered in *cis*. Our results are consistent with a model where longer, lowly expressed, highly connected, and highly modified proteins are “buffered” from phosphorylation variation in *cis*.

## DISCUSSION

### Technical assessment and recommendation for study design

In our study we applied an LC-MS/MS approach capable of deeply and reproducibly profiling the phosphoproteome to capture germline genetically-driven differential phosphorylation. We applied the same SILAC LCL standard [81] that we used before [34, 49] in order to facilitate an integrative analysis. Critically, our application of a common SILAC standard allowed us to assess phosphorylation variation while controlling for variation in protein expression. This property of our study design also allowed us to address the systematic error produced from our multi-fraction technical workup (**Supplemental Figure 1**). For our cell system and experimental approach, the magnitude of the donor associated variance component was on average much greater than either nested variance component, resulting in reasonable power to detect extensive differential phosphorylation between individuals (**Figure 2**). As all our LCLs samples were of similar passage and were cultured in practically identical conditions, we assume that environmental impact on differences between individuals is minimal. In other words, we assume that inter-individual variation in phosphorylation in our study stems from genetically-driven differences.

Given our inference that the genetic background drives most of the observed phosphopeptide variation, a less precise but also less laborious and time-consuming approach may actually be feasible for future studies. Recent reports of label-free LC-MS/MS phosphoproteomic approaches demonstrate greatly improved phosphoproteome sampling depth over previous label-free methods [14, 15]. While a label-free approach is unlikely to achieve the same phosphoproteome coverage as our multi-fraction protocol, label-free approaches have other benefits, such as an independence from the requirement to quantify peptides from a common standard. This independence would enable the quantification of phosphopeptides that are not found in a common standard but may be prevalent in other samples. Given that our results can inform future LC-MS/MS methodology choices, we performed simulations to assess the impact technical variance has on power (see methods). It is important to note that while this power analysis was performed without explicit regard to variation in DNA sequence, the effect sizes used are directly relevant for future phosphorylation QTL mapping studies where the aim is to identify relationships between coding variants and phosphorylation levels in *cis.* We found (**Supplemental Figure 8)** that for our design (2 technical replicates), the power lost by increasing the technical variance up to 5-fold is almost completely compensated for by doubling the number of technical replicates. For our system (LCLs), this would be a welcome trade considering that this translates into less than half our currently required protein input and 1/8 of the instrument time (assuming 1 mg input and 1.5 hr gradients as in [14]).

### Functional associations with phosphopeptide variation

We uncovered multiple intriguing correlations between putatively genetically-driven phosphopeptide variation and functional annotations of polymorphisms and proteins. Of note is the apparently greater impact coding variants (especially those mapping to domains or those predicted to have functional consequences) have on phosphopeptide variation in *cis* relative to variants known to impact protein expression levels (**Table 2**). This observation implies that information relay via phosphorylation is more robust to variation in substrate protein levels than variation in substrate protein structure. This dichotomy may also portend a lack of concordance between pQTLs and phosQTLs (similar to that recently reported between eQTL and pQTL [34]) and relatively greater concordance between splicing QTLs and phosQTLs [40]. Future work employing additional samples is required to explore this property further.

We also uncovered novel aspects of phosphopeptide variation. For example, we observed an enrichment of phosphopeptides that map to functional protein segments (domains or annotated motifs) amongst highly variable phosphopeptides. We also consistently observed that PRPs such as kinases and phosphatases do not, on average, possess more variable phosphopeptides. From a systems perspective, we uncovered that the interaction count, PTM count, and disorder content of a protein correlate negatively with phosphopeptide variation in *cis*. Increased levels of these systems-relevant annotations are characteristic of longer, lowly expressed and tightly regulated proteins that are of amplified importance within interaction networks [74, 79, 80]. Intriguingly, the decreased variability of phosphopeptides mapping to such proteins may protect the cell from adopting unfavorable signaling states. We also observed increased inter-individual variability for phosphopeptides mapping to highly expressed proteins. This may result from random encounters with kinases and phosphatases and would therefore imply a lack of function [69]. Indeed, recent reports have found that highly expressed proteins are enriched in low stoichiometry phosphorylation sites with conservation rates similar to those of their non-phosphorylated counterparts [69] and estimate that 80% of cellular ATP is consumed by only 20% of the (putatively functional) phosphorylation sites [5]. Future work will benefit from the application of these insights to prioritize phosphorylation events for further mechanistic characterization.

### Conclusions and Future Directions

We characterized inter-individual variation in PTM levels at substantial sampling depth with genotyped human cell lines. We provided evidence that variants affecting either protein structure or protein expression are associated with inter-individual phosphorylation variation. Our observations suggest that protein length, connectivity, and/or expression level may serve as a functional buffer against inter-individual phosphorylation variation. The generality of these results with respect to cell type, stimulation conditions, and sample size is currently unknown and requires further study. Lastly, our study demonstrates that current phosphoproteomic LC-MS/MS protocols are sufficient to capture germline driven PTM variation and provides a context for further technical development toward this end.

## MATERIALS AND METHODS

### Cell culturing and SILAC labeling

Epstein-Barr virus (EBV)-transformed lymphoblastoid cell lines (LCLs) derived from Yoruba individuals in Ibadan, Nigeria (YRI from Coriell, NIGMS Human Genetic Cell Repository) were cultured under identical conditions of 37°C and 5% CO_2_. Each of the three lines (GM18486, GM19160, and GM18862) were grown in Lys/Arg depleted RPMI and 15% dialyzed FBS supplemented with 2 mM L-glutamate, 100 IU/ml penicillin, 100 μg/ml streptomycin and L-^13^C_6_^15^N_4_-Arg (Arg-10) and L-^13^C_6_^15^N_2_-Lys (Lys-8) (Cambridge Isotopes, Andover, USA). Each line was cultured to ~200 × 10^6^ cells over at least six doublings. Culture replicates were awoken from the same frozen pellet and cultured in parallel. Label incorporation was verified by analyzing the protein lysate from the labeled LCLs alone by high-resolution LC-MS/MS. The internal (unlabeled) standard line (GM19238) was expanded to 20 × 10^9^ cells in roller bottles using RPMI media with 15% FBS and 2 mM L-glutamate by the Coriell Institute for Medical Research.

### Genotype data

The genotypes for the three YRI individuals were collected as part of the International HapMap Project [42]. Additional SNPs from the 1000 Genomes Phase1 integrated version 3 reference panel [82] were imputed using IMPUTE2 [83], as previously described [36]. SnpEff [84] and SnpSift [85] were used to identify all SNPs which had an effect on the amino acid sequence of a protein annotated in Ensembl GRCh37 release 75. PolyPhen2 [58] predictions were sourced from dbNSFP [86] via SNPSift. A variant was included for analysis if it was observed in at least one allele in at least one of the LCLs except in the case where each LCL has the variant with the same genotype.

### Quantitative, high-resolution mass spectrometry

Cell pellets were washed twice with 500μL 25mM ammonium bicarbonate and centrifuged at 5000 × g for 2 minutes. Washes were discarded. Cell lysis was performed with 3mL of urea lysis buffer (8M urea, 50mM Tris.HCl pH8, 100mM NaCl) using 3 applications of a Qsonica Q125 sonic probe with a 30 second pulse and 80% amplitude. The cell lysate was centrifuged at 10,000g for 10 minutes at 25°C. The protein concentration of the cleared lysate was determined with a Qubit protein assay (Invitrogen). For each experiment 4.5mg of light and 4.5mg heavy protein were combined and digested with the following protocol: Reduction with 10mM dithiothreitol at 25°C for 30 minutes followed by alkylation with 20mM iodoacetamide at 25°C for 45 minutes. Proteins were digested with 200μg sequencing grade trypsin (Promega) at 37°C overnight. The final digest volume was 25mL adjusted with 25mM ammonium bicarbonate. The digestion was cooled to room temperature and terminated with 5μL of formic acid. The digest was centrifuged at 10,000g for 10 minutes. Peptides were desalted with 500mg Sep-Pak (Waters) and dried using vacuum centrifugation in a SpeedVac. Dried peptides were dissolved in 7 mM KH_2_PO_4_, pH 2.65, 30% ACN and protein quantitation performed with a 280nm protein assay. Peptides were fractionated on an Agilent 1100 equipped with a 500 uL sample loop operating at 2 mL/min, detector set at 220-280 nm wavelengths. 5mg of peptide was loaded on polySULFOETHYL A, 4.6 mm ID × 200 mm length, 5 um particle size, 200 Å pore size (polyLC, from the Nest group). A total of 48 fractions were collected at 1 min intervals. In batches of 3, adjacent SCX fractions were pooled and processed by solid phase extraction (SPE) using a Waters SEP-PAK 50mg C18 cartridge per the vendor protocol and dried overnight in a lyophilizer. Phosphopeptides were enriched using Titansphere TiO_2_ tips from GL sciences using the vendor protocol. Phosphopeptides were eluted from the tips using two eluents: 50μL 5% NH_4_OH in water and 50μL 5% Pyrrolidine in water. The eluents were combined and neutralized with 50% acetic acid and dried. Dry samples were reconstituted in 100μL 0.1% trifluoroacetic (TFA) acid. Each enriched sample was desalted using a Stage Tip (ThermoFisher P/N SP301) per the vendor protocol. Peptides were dried and reconstituted in 70μL of 0.1% TFA prior to analysis.

Half of each enriched sample was analyzed by nano LC-MS/MS with a Waters NanoAcquity HPLC system interfaced to a ThermoFisher Q Exactive mass spectrometer. Peptides were loaded on a trapping column and eluted over a 75μm analytical column at 350nL/min using a 2hr reverse phase gradient; both columns were packed with Jupiter Proteo resin (Phenomenex). The injection volume was 30μL. The mass spectrometer was operated in data-dependent mode, with the Orbitrap operating at 60,000 FWHM and 17,500 FWHM for MS and MS/MS respectively. The fifteen most abundant ions were selected for MS/MS.

### Computational analysis of mass spectrometry data

MS data was analyzed with MaxQuant [46] 1.5.0.30 and the Adromeda [47] search engine. Proteins were identified using a protein sequence database containing 35,585 consensus coding sequences (CCDS) [87] translated from GRCh37/hg19 gene models using Ensembl release 75 annotations. Only translations of genes/transcripts of status ‘known’ or ‘novel’ and biotype ‘protein coding’ were used. Each sequence has a unique protein identifier (ENSP ID) that allows for mapping between gene and transcript IDs. Carbamidomethylation of cysteine was allowed as a fixed modification. N-terminal acetylation and oxidation of methionine as variable modifications were included for all searches, while phosphorylation of S/T/Y was included for the phosphorylation data. Up to three missed cleavages were allowed for phosphoproteomic data and two missed cleavages were allowed for proteomic data. A ‘first search’ tolerance of 40 ppm with a score threshold of 75 was used for time-dependent recalibration followed by a main search MS1 tolerance of 6 ppm and an MS2 tolerance of 20 ppm. The ‘re-quantify’ option was used to aid the assignment of isotopic patterns to labeling pairs. The ‘match between runs’ option was enabled to match identifications across samples using a matching time window of 42 seconds and an alignment time window of 30 min for phosphoproteomic data and 20 min for proteomic data. Peptide and protein false discovery rates were set to 1%. Quantitative analysis of phosphorylated peptides was limited to ‘class 1’ sites with median localization probability of .99 and minimum localization probability of .75. Protein group quantifications were taken as the median log_2_(sample/standard) ratio for all groups containing at least two independent unique or ‘razor’ peptide quantifications (including multiple measurements of the same peptide in different fractions) without a modified peptide counterpart. For the purposes of protein level normalization, a custom R script was used to assign phosphopeptides to protein groups based on the presence of the peptide within the sequences of the protein group proteins. In the rare cases where multiple protein groups contain proteins that match a phosphopeptide, the protein group with the most peptide identifications is assigned to the phoshpopeptide. For enrichment analyses, members of the protein group were parsed such that each member contained the given phosphopeptide sequence.

### Normalization and batch correction

Log-transformed phosphopeptide SILAC ratios were normalized in two steps. First, within each sample, we centered the log intensity ratios by subtracting the sample-specific median log intensity ratio. Second, we applied quantile normalization to account for between sample variation in log intensity ratios with the *limma* [88] function ‘normalizeQuantiles’. We used ComBat [48] to estimate and adjust for sample variation in the log intensity ratios attributable to processing date (with the *swamp* [89] function ‘combat’).

### Variance component analysis

To assess the relative contributions of individual (donor identify), culture (technical replication of the cell culture), and technical workup (protein sample processing and MS workflow) variation to the observed logged SILAC intensity ratios of phosphopeptides, we fit a linear mixed model for each phosphopeptide on the quantile-normalized, batch effect-corrected data. The model estimates variance components due to the random effect of individual, culture, and technical workup as follows:

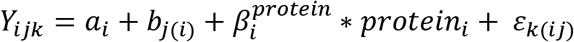

where *Y*_*ijk*_ denotes the observed logged intensity ratio of individual *i*, derived from the *k*-th technical workup of the *j*-th culture replicate, with *i* = GM18486, GM19160, GM18862, *j* =1, 2, *k* =1, 2. *a*_*i*_ denotes random effects of individual, *b*_*j*(*i*)_ estimates the random effect of *j*-th culture replicate for individual *i*, and *ε*_*k*(*ij*)_ denotes the random effect of *k*-th technical workup in *j*-th culture replicate for individual *i*. Protein expression levels 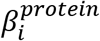 in individual *i* are included as fixed covariates to account for the confounding effect of variation in protein expression levels. The protein expression measurements for each phosphopeptide were derived from SILAC protein expression ratios previously reported in an MS dataset collected from 60 Yoruba LCLs [49]. These protein data were processed as described under our Results section entitled *Characterization of genetically-driven differential phosphorylation*. The random effects of individual sample *a*_*i*_, culture replicate *b*_*j*(*i*)_, and technical workup *ε*_*k*(*ij*)_ are assumed to be independent and follow normal distributions with zero mean and variance components 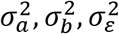, respectively. The R package *MCMCglmm* [90] was used to estimate the variance components associated with the random effects. A similar analysis was performed for **Supplemental Figure 4** using normalized phosphopeptide data and processing date batch as a covariate.

### Differential phosphorylation analysis

To quantify individual differences for each phosphopeptide in the observed logged SILAC intensity ratios, we fit a linear mixed effect model on the quantile-normalized, batch effect-corrected data: including individual as a fixed effect, culture replication as a random effect and logged protein SILAC intensity ratios as a fixed covariate. Our approach is based on *limma* – a popular linear-model based approach for differential abundance analysis in genome-wide expression studies. In our model, we also include weights for each phosphopeptide to account for the relationship between the model residuals of SILAC intensity ratios and the average log intensity of the phosphopeptides. Specifically, the model residuals of SILAC intensity ratios are negatively correlated with log intensity of the phosphopeptides. We computed observation-level weights of the model residuals using the *voom* approach [91]. A similar approach was used in previous work on modeling differential abundance in label-free LC-MS/MS proteomic experiments [92].

Furthermore, we explicitly accounted for noise in the protein level estimates by a commonly used strategy for maximizing the accuracy of molecular measurements in regulatory eQTL studies [34] (see Results section for more details). Briefly, we serially regressed PCs from the full protein data matrix derived from 60 LCLs [34] and identified pQTLs from the resulting residual matrix. The residual matrix with the first 13 PCs regressed produced the maximum pQTL count and was therefore employed here. The Benjamini and Hochberg [93] procedure was used to compute false discovery rates (FDR) via the ‘p.adjust’ function from the R package *stats* [94]. Significant individual variation in the phosphopeptide intensity ratio was identified at 5% FDR.

### Power analysis

We investigated the power of our differential phosphorylation analysis to detect significant individual variation. Specifically, we estimated the number of technical replicates required (per culture replicate) to reach 80% power, given varying levels of sampling noise from technical workup. The power calculation proceeds as follows:

1. Identify phosphopeptides with significant individual variation, and among these, choose the one phosphopeptide with the largest p-value and compute its effect size (F-statistic).
2. Based on the choice of phosphopeptide, extract all parameter estimates in the differential phosphorylation analysis, including the effect sizes of individual variation (F-statistic) and protein expression covariate, and the variance components of culture replicate and technical workup.
3. Simulate 1,000 peptides under the model assumptions of differential phosphorylation analysis. Use parameter estimates from Step 2. Fix the effect sizes of individual variation and protein expression levels, and the variance component of culture replicate. Vary the number of technical replicates and variance components of technical workup.
4. Given the settings in Step 2, we compute power as the probability of detecting significant individual variation in each simulation at FDR 5%.

### Protein, domain and phosphosite annotation

Pfam [95] domain assignment and boundary definition was accomplished using InterProScan [96] and Ensembl 75 CCDS FASTAs. Kinases, phosphatases, and modular phosphopeptide binding domains with the following Pfam family identifiers were considered functionally relevant for phosphorylation mediated signaling: PF00498, PF01846, PF03166, PF10401, PF00244, PF00533, PF00400, PF00659, PF00397, PF00782, PF06602, PF04273, PF14566, PF14671, PF04179, PF05706, PF00069, PF01636, PF03109, PF03881, PF06293, PF01163, PF01633, PF10707, PF06176, PF02958, PF04655, PF10009, PF12260, PF16474, PF07914, PF14531, PF06734, PF05445, PF07387. Gene Ontology [97] IDs were sourced using biomaRt [98]. PTM site datasets were sourced from PhosphoSitePlus [99] on 9/8/15. Human physical protein-protein interaction data was sourced from BioGRID [100] v3.4.127 on 8/25/15. Eukaryotic Linear Motif (ELM) [61] instances were sourced on 10/7/15 and mapped to proteins and phosphosites using custom R scripts. Kinase motifs were sourced from the Human Protein Reference Database (HPRD) [101] via MaxQuant’s Perseus module. Protein level disorder was predicted using the RAPID [102] algorithm and webserver on 8/28/15. Amino acid disorder was predicted with IUPred [103]. Scores ≥ .5 were considered disordered.

### Enrichment analyses

We assessed gene ontology enrichment on phosphopeptides categorized as differentially expressed (5 % FDR) across each contrast using one-sided Fisher’s exact tests. Adjusted p-values were derived using the approach of Benjamini and Hochberg [93] via the ‘p.adjust’ function from the ‘stats’ R package [94]. Distributions of nominal p-values derived from *limma* omnibus F-tests were used to assess the enrichment of annotations as outlined previously [34]. For each test, we calculated Spearman’s correlations between a vector of negative log-transformed *limma* F-test p-values and a binary vector designating assignment of the protein containing the phosphopeptide, or the phosphopeptide itself to an annotation. The p-value for the Spearman’s correlation was computed with the R function ‘cor.test’ with the option ‘exact = FALSE’. Amino acid position specific enrichments were produced with ‘pLogo’ [104].

### Code and data availability

The custom R [94] scripts used in this study are available from https://github.com/bengalengel/Phospilot. The mass spectrometry proteomics data have been deposited to the ProteomeXchange Consortium via the PRIDE [105] partner repository with the dataset identifier PXD008002. SILAC protein estimates are available from proteomeXchange; identifier PXD001406. pQTL data are available from http://www.sciencemag.org/content/suppl/2014/12/17/science.1260793.DC1/1260793_DatafileS1.xlsx. Genotype data are available from http://eqtl.uchicago.edu/jointLCL/genotypesYRI.gen.txt.gz

## Acknowledgements

This work was funded by NIH grant HL092206. BWE was supported in part by a Ruth L. Kirschstein NRSA fellowship (F32GM116390-01). CJH was supported by the grant U01CA198933 from the NIH BD2K program.

## Author contributions

B.W.E. and Y.G. conceived and designed the study. B.W.E. and M.F. performed experiments. B.W.E., C.J.H., J.D.B., Y.F., and Y.G. analyzed and interpreted the data. B.W.E., C.J.H., J.D.B., and Y.G. drafted or revised the article.

## References

1. Ubersax, J.A. and J.E. Ferrell, Jr., Mechanisms of specificity in protein phosphorylation. Nat Rev Mol Cell Biol, 2007. 8(7): p. 530–41.

2. Lim, W., B. Mayer, and T. Pawson, Cell Signaling: Principles and mechanisms. 1 ed. 2014, New York, New York: Garland Science. 412.

3. Deribe, Y.L., T. Pawson, and I. Dikic, Post-translational modifications in signal integration. Nat Struct Mol Biol, 2010. 17(6): p. 666–672.

4. Manning, G., et al., The protein kinase complement of the human genome. Science, 2002. 298(5600): p. 1912–34.

5. Sharma, K., et al., Ultradeep human phosphoproteome reveals a distinct regulatory nature of Tyr and Ser/Thr-based signaling. Cell Rep, 2014. 8(5): p. 1583–94.

6. Tartaglia, M., et al., Mutations in PTPN11, encoding the protein tyrosine phosphatase SHP-2, cause Noonan syndrome. Nat Genet, 2001. 29(4): p. 465–8.

7. Blume-Jensen, P. and T. Hunter, Oncogenic kinase signalling. Nature, 2001. 411(6835): p. 355–65.

8. Tan, C.S., et al., Comparative analysis reveals conserved protein phosphorylation networks implicated in multiple diseases. Sci Signal, 2009. 2(81): p. ra39.

9. Hanahan, D. and R.A. Weinberg, Hallmarks of cancer: the next generation. Cell, 2011. 144(5): p. 646–74.

10. Bensimon, A., A.J. Heck, and R. Aebersold, Mass spectrometry-based proteomics and network biology. Annu Rev Biochem, 2012. 81: p. 379–405.

11. Pe’er, D. and N. Hacohen, Principles and strategies for developing network models in cancer. Cell, 2011. 144(6): p. 864–73.

12. Linding, R., et al., Systematic discovery of in vivo phosphorylation networks. Cell, 2007. 129(7): p. 1415–26.

13. Ellis, M.J., et al., Connecting genomic alterations to cancer biology with proteomics: the NCI Clinical Proteomic Tumor Analysis Consortium. Cancer Discov, 2013. 3(10): p. 1108–12.

14. de Graaf, E.L., et al., Single-step enrichment by Ti4+-IMAC and label-free quantitation enables in-depth monitoring of phosphorylation dynamics with high reproducibility and temporal resolution. Mol Cell Proteomics, 2014. 13(9): p. 2426–34.

15. Humphrey, S.J., S.B. Azimifar, and M. Mann, High-throughput phosphoproteomics reveals in vivo insulin signaling dynamics. Nat Biotechnol, 2015. 33(9): p. 990–5.

16. Creixell, P., et al., Kinome-wide Decoding of Network-Attacking Mutations Rewiring Cancer Signaling. Cell. 163(1): p. 202–217.

17. Rotival, M., et al., Integrating phosphoproteome and transcriptome reveals new determinants of macrophage multinucleation. Mol Cell Proteomics, 2015. 14(3): p. 484–98.

18. Phanstiel, D.H., et al., Proteomic and phosphoproteomic comparison of human ES and iPS cells. Nat Meth, 2011. 8(10): p. 821–827.

19. Mertins, P., et al., Proteogenomics connects somatic mutations to signalling in breast cancer. Nature, 2016. 534(7605): p. 55–62.

20. Zhang, H., et al., Integrated Proteogenomic Characterization of Human High-Grade Serous Ovarian Cancer. Cell, 2016. 166(3): p. 755–765.

21. Miraldi, E.R., et al., Molecular network analysis of phosphotyrosine and lipid metabolism in hepatic PTP1b deletion mice. Integr Biol (Camb), 2013. 5(7): p. 940–63.

22. Boekhorst, J., et al., Comparative phosphoproteomics reveals evolutionary and functional conservation of phosphorylation across eukaryotes. Genome Biol, 2008. 9(10): p. R144.

23. Freschi, L., M. Osseni, and C.R. Landry, Functional Divergence and Evolutionary Turnover in Mammalian Phosphoproteomes. PLOS Genetics, 2014. 10(1): p. e1004062.

24. Reimand, J., O. Wagih, and G.D. Bader, Evolutionary Constraint and Disease Associations of Post-Translational Modification Sites in Human Genomes. PLOS Genetics, 2015. 11(1): p. e1004919.

25. Radivojac, P., et al., Gain and loss of phosphorylation sites in human cancer. Bioinformatics, 2008. 24(16): p. i241–7.

26. Reimand, J. and G.D. Bader, Systematic analysis of somatic mutations in phosphorylation signaling predicts novel cancer drivers. Mol Syst Biol, 2013. 9: p. 637.

27. Wagih, O., J. Reimand, and G.D. Bader, MIMP: predicting the impact of mutations on kinase-substrate phosphorylation. Nat Meth, 2015. 12(6): p. 531–533.

28. Creixell, P., et al., Kinome-wide decoding of network-attacking mutations rewiring cancer signaling. Cell, 2015. 163(1): p. 202–17.

29. Mackay, T.F.C., E.A. Stone, and J.F. Ayroles, The genetics of quantitative traits: challenges and prospects. Nat Rev Genet, 2009. 10(8): p. 565–577.

30. Foss, E.J., et al., Genetic basis of proteome variation in yeast. Nat Genet, 2007. 39(11): p. 1369–75.

31. Pickrell, J.K., et al., Understanding mechanisms underlying human gene expression variation with RNA sequencing. Nature, 2010. 464(7289): p. 768–72.

32. Consortium, G.T., The Genotype-Tissue Expression (GTEx) project. Nat Genet, 2013. 45(6): p. 580–5.

33. Wu, L., et al., Variation and genetic control of protein abundance in humans. Nature, 2013. 499(7456): p. 79–82.

34. Battle, A., et al., Genomic variation. Impact of regulatory variation from RNA to protein. Science, 2015. 347(6222): p. 664–7.

35. Banovich, N.E., et al., Methylation QTLs Are Associated with Coordinated Changes in Transcription Factor Binding, Histone Modifications, and Gene Expression Levels. PLoS Genet, 2014. 10(9): p. e1004663.

36. McVicker, G., et al., Identification of genetic variants that affect histone modifications in human cells. Science, 2013. 342(6159): p. 747–9.

37. Degner, J.F., et al., DNase I sensitivity QTLs are a major determinant of human expression variation. Nature, 2012. 482(7385): p. 390–4.

38. Veyrieras, J.-B., et al., Exon-Specific QTLs Skew the Inferred Distribution of Expression QTLs Detected Using Gene Expression Array Data. PLoS ONE, 2012. 7(2): p. e30629.

39. Veyrieras, J.B., et al., High-resolution mapping of expression-QTLs yields insight into human gene regulation. PLoS Genet, 2008. 4(10): p. e1000214.

40. Li, Y.I., et al., RNA splicing is a primary link between genetic variation and disease. Science, 2016. 352(6285): p. 600–4.

41. The International HapMap Project. Nature, 2003. 426(6968): p. 789–796.

42. Frazer, K.A., et al., A second generation human haplotype map of over 3.1 million SNPs. Nature, 2007. 449(7164): p. 851–61.

43. Ong, S.E., et al., Stable isotope labeling by amino acids in cell culture, SILAC, as a simple and accurate approach to expression proteomics. Mol Cell Proteomics, 2002. 1(5): p. 376–86.

44. Villen, J. and S.P. Gygi, The SCX/IMAC enrichment approach for global phosphorylation analysis by mass spectrometry. Nat Protoc, 2008. 3(10): p. 1630–8.

45. Michalski, A., et al., Mass spectrometry-based proteomics using Q Exactive, a high-performance benchtop quadrupole Orbitrap mass spectrometer. Mol Cell Proteomics, 2011. 10(9): p. M111.011015.

46. Cox, J. and M. Mann, MaxQuant enables high peptide identification rates, individualized p.p.b.–range mass accuracies and proteome-wide protein quantification. Nat Biotechnol, 2008. 26(12): p. 1367–72.

47. Cox, J., et al., Andromeda: a peptide search engine integrated into the MaxQuant environment. J Proteome Res, 2011. 10(4): p. 1794–805.

48. Johnson, W.E., C. Li, and A. Rabinovic, Adjusting batch effects in microarray expression data using empirical Bayes methods. Biostatistics, 2007. 8(1): p. 118–27.

49. Khan, Z., et al., Primate transcript and protein expression levels evolve under compensatory selection pressures. Science, 2013. 342(6162): p. 1100–4.

50. Choy, E., et al., Genetic Analysis of Human Traits In Vitro: Drug Response and Gene Expression in Lymphoblastoid Cell Lines. PLOS Genetics, 2008. 4(11): p. e1000287.

51. Parker, C.C., et al., Genome-wide association study of behavioral, physiological and gene expression traits in outbred CFW mice. Nat Genet, 2016. 48(8): p. 919–926.

52. Ponting, C.P. and R.R. Russell, The natural history of protein domains. Annu Rev Biophys Biomol Struct, 2002. 31: p. 45–71.

53. Jin, J., et al., Eukaryotic protein domains as functional units of cellular evolution. Sci Signal, 2009. 2(98): p. ra76.

54. Hornbeck, P.V., et al., PhosphoSitePlus, 2014: mutations, PTMs and recalibrations. Nucleic Acids Res, 2015. 43(Database issue): p. D512–20.

55. Davey, N.E., et al., Attributes of short linear motifs. Mol Biosyst, 2012. 8(1): p. 268–81.

56. Babu, M.M., et al., Intrinsically disordered proteins: regulation and disease. Curr Opin Struct Biol, 2011. 21(3): p. 432–40.

57. Perkins, J.R., et al., Transient protein-protein interactions: structural, functional, and network properties. Structure, 2010. 18(10): p. 1233–43.

58. Adzhubei, I.A., et al., A method and server for predicting damaging missense mutations. Nat Methods, 2010. 7(4): p. 248–9.

59. Seet, B.T., et al., Reading protein modifications with interaction domains. Nat Rev Mol Cell Biol, 2006. 7(7): p. 473–83.

60. Hunter, T., Protein kinases and phosphatases: the yin and yang of protein phosphorylation and signaling. Cell, 1995. 80(2): p. 225–36.

61. Puntervoll, P., et al., ELM server: A new resource for investigating short functional sites in modular eukaryotic proteins. Nucleic Acids Res, 2003. 31(13): p. 3625–30.

62. Zhou, S., et al., SH2 domains recognize specific phosphopeptide sequences. Cell. 72(5): p. 767–778.

63. Songyang, Z., et al., Use of an oriented peptide library to determine the optimal substrates of protein kinases. Curr Biol, 1994. 4(11): p. 973–82.

64. Obenauer, J.C., L.C. Cantley, and M.B. Yaffe, Scansite 2.0: Proteome-wide prediction of cell signaling interactions using short sequence motifs. Nucleic Acids Res, 2003. 31(13): p. 3635–41.

65. Miller, M.L., et al., Linear motif atlas for phosphorylation-dependent signaling. Sci Signal, 2008. 1(35): p. ra2.

66. Engelmann, B.W., et al., The Development and Application of a Quantitative Peptide Microarray Based Approach to Protein Interaction Domain Specificity Space. Molecular & Cellular Proteomics, 2014. 13(12): p. 3647–3662.

67. Stein, A. and P. Aloy, Contextual specificity in peptide-mediated protein interactions. PLoS One, 2008. 3(7): p. e2524.

68. Landry, C.R., E.D. Levy, and S.W. Michnick, Weak functional constraints on phosphoproteomes. Trends Genet, 2009. 25(5): p. 193–7.

69. Levy, E.D., S.W. Michnick, and C.R. Landry, Protein abundance is key to distinguish promiscuous from functional phosphorylation based on evolutionary information. Philos Trans R Soc Lond B Biol Sci, 2012. 367(1602): p. 2594–606.

70. Gnad, F., et al., PHOSIDA (phosphorylation site database): management, structural and evolutionary investigation, and prediction of phosphosites. Genome Biol, 2007. 8(11): p. R250.

71. Chen, S.C., F.C. Chen, and W.H. Li, Phosphorylated and nonphosphorylated serine and threonine residues evolve at different rates in mammals. Mol Biol Evol, 2010. 27(11): p. 2548–54.

72. Wang, Z., et al., Evolution of protein phosphorylation for distinct functional modules in vertebrate genomes. Mol Biol Evol, 2011. 28(3): p. 1131–40.

73. Fuxreiter, M., P. Tompa, and I. Simon, Local structural disorder imparts plasticity on linear motifs. Bioinformatics, 2007. 23(8): p. 950–6.

74. Jeong, H., et al., Lethality and centrality in protein networks. Nature, 2001. 411.

75. Haynes, C., et al., Intrinsic disorder is a common feature of hub proteins from four eukaryotic interactomes. PLoS Comput Biol, 2006. 2(8): p. e100.

76. Xie, H., et al., Functional anthology of intrinsic disorder. 3. Ligands, post-translational modifications, and diseases associated with intrinsically disordered proteins. J Proteome Res, 2007. 6(5): p. 1917–32.

77. Pawson, T. and P. Nash, Assembly of cell regulatory systems through protein interaction domains. Science, 2003. 300(5618): p. 445–52.

78. Ekman, D., et al., What properties characterize the hub proteins of the protein-protein interaction network of Saccharomyces cerevisiae? Genome Biology, 2006. 7(6): p. 1–13.

79. Vavouri, T., et al., Intrinsic Protein Disorder and Interaction Promiscuity Are Widely Associated with Dosage Sensitivity. Cell. 138(1): p. 198–208.

80. Gsponer, J., et al., Tight regulation of unstructured proteins: from transcript synthesis to protein degradation. Science, 2008. 322(5906): p. 1365–8.

81. Geiger, T., et al., Super-SILAC mix for quantitative proteomics of human tumor tissue. Nat Methods, 2010. 7(5): p. 383–5.

82. Abecasis, G.R., et al., An integrated map of genetic variation from 1,092 human genomes. Nature, 2012. 491(7422): p. 56–65.

83. Howie, B.N., P. Donnelly, and J. Marchini, A flexible and accurate genotype imputation method for the next generation of genome-wide association studies. PLoS Genet, 2009. 5(6): p. e1000529.

84. Cingolani, P., et al., A program for annotating and predicting the effects of single nucleotide polymorphisms, SnpEff: SNPs in the genome of Drosophila melanogaster strain w1118; iso-2; iso-3. Fly (Austin), 2012. 6(2): p. 80–92.

85. Cingolani, P., et al., Using Drosophila melanogaster as a Model for Genotoxic Chemical Mutational Studies with a New Program, SnpSift. Front Genet, 2012. 3: p. 35.

86. Liu, X., X. Jian, and E. Boerwinkle, dbNSFP: A lightweight database of human nonsynonymous SNPs and their functional predictions. Human Mutation, 2011. 32(8): p. 894–899.

87. Pruitt, K.D., et al., The consensus coding sequence (CCDS) project: Identifying a common protein-coding gene set for the human and mouse genomes. Genome Res, 2009. 19(7): p. 1316–23.

88. Ritchie, M.E., et al., limma powers differential expression analyses for RNA-sequencing and microarray studies. Nucleic Acids Res, 2015. 43(7): p. e47.

89. Lauss, M., swamp: Visualization, analysis and adjustment of high-dimensional data in respect to sample annotations. 2013.

90. Hadfield, J.D., MCMC Methods for Multi-Response Generalized Linear Mixed Models: The {MCMCglmm} {R} Package. Journal of Statistical Software, 2010. 33(2): p. 1–22.

91. Law, C.W., et al., voom: Precision weights unlock linear model analysis tools for RNA-seq read counts. Genome Biol, 2014. 15(2): p. R29.

92. Clough, T., et al., Statistical protein quantification and significance analysis in label-free LC-MS experiments with complex designs. BMC Bioinformatics, 2012. 13 Suppl 16: p. S6.

93. Benjamini, Y. and Y. Hochberg, Controlling the False Discovery Rate: A Practical and Powerful Approach to Multiple Testing. Journal of the Royal Statistical Society. Series B (Methodological), 1995. 57(1): p. 289–300.

94. R Development Core Team, R: A Language and Environment for Statistical Computing. 2015, R Foundation for Statistical Computing: Vienna, Austria.

95. Sonnhammer, E.L., et al., Pfam: multiple sequence alignments and HMM-profiles of protein domains. Nucleic Acids Res, 1998. 26(1): p. 320–2.

96. Quevillon, E., et al., InterProScan: protein domains identifier. Nucleic Acids Research, 2005. 33(suppl 2): p. W116–W120.

97. Ashburner, M., et al., Gene ontology: tool for the unification of biology. The Gene Ontology Consortium. Nat Genet, 2000. 25(1): p. 25–9.

98. Durinck, S., et al., Mapping identifiers for the integration of genomic datasets with the R/Bioconductor package biomaRt. Nat Protoc, 2009. 4(8): p. 1184–91.

99. Hornbeck, P.V., et al., PhosphoSitePlus: a comprehensive resource for investigating the structure and function of experimentally determined post-translational modifications in man and mouse. Nucleic Acids Res, 2012. 40(Database issue): p. D261–70.

100. Stark, C., et al., BioGRID: a general repository for interaction datasets. Nucleic Acids Res, 2006. 34(Database issue): p. D535–9.

101. Prasad, T.S., K. Kandasamy, and A. Pandey, Human Protein Reference Database and Human Proteinpedia as discovery tools for systems biology. Methods Mol Biol, 2009. 577: p. 67–79.

102. Yan, J., et al., RAPID: fast and accurate sequence-based prediction of intrinsic disorder content on proteomic scale. Biochim Biophys Acta, 2013. 1834(8): p. 1671–80.

103. Dosztányi, Z., et al., IUPred: web server for the prediction of intrinsically unstructured regions of proteins based on estimated energy content. Bioinformatics, 2005. 21(16): p. 3433–3434.

104. O’Shea, J.P., et al., pLogo: a probabilistic approach to visualizing sequence motifs. Nat Methods, 2013. 10(12): p. 1211–2.

105. Vizcaino, J.A., et al., 2016 update of the PRIDE database and its related tools. Nucleic Acids Res, 2016. 44(22): p. 11033.

